# Selecting cacao clones with high productivity potential and tolerance to Black Pod Rot (BPR) and Frosty Pod Rot (FPR)

**DOI:** 10.1101/2024.10.31.621355

**Authors:** Marlon Enrique López, Oscar Arnulfo Ramírez, Aroldo Dubón, Javier Díaz

**Affiliations:** Cacao and Agroforestry Program, Honduran Foundation for Agricultural Research (FHIA), La Lima Cortés Honduras

**Keywords:** Disease and Production Index, Moniliophthora, Phytophthora, Cacao Production, FHIA

## Abstract

Cacao (*Theobroma cacao*) is affected annually by Frosty Pod Rot (FPR) (*Moniliopthora roreri*) and Black Pod Rot (BPR) (*Phytophthora* sp). The loss caused by both diseases is threatening cacao production worldwide. Therefore, cacao breeding programs focus on developing new cacao clones with high productivity potential and disease resistance. However, this challenge is not easy to achieve due to the long time required for selection, the influence of environmental conditions on the severity of the disease, and the avoidance of chemical control of diseases. Therefore, genetic resistance should be the best option for selecting and releasing new cacao clones to the farmers. In this study, 40 cacao clones, 20 from CATIE and 20 from FHIA breeding programs were evaluated from 2013 to 2017. Three criteria were used to select clones: Yield, Percentage Disease Pod (PDP), and Disease and Production Index (DPI). Results indicated that depending on the cacao breeding program objectives, it is possible to use those criteria to select new cacao clones that are highly productive and resistant to diseases because cacao clones with high productivity are not always the most resistant to diseases and vice versa; however, a combination of both criteria can be used to select cacao clones with high productivity potential and resistance to FPR and BPR.

## 1. INTRODUCTION

Cacao production extends throughout the world and is threatened by pests and diseases; approximately one-third of global production is lost annually (Marelli et al., 2019). Four diseases account for the most significant losses worldwide: Black Pod Rot (BPR), caused by four *Phytophthora* spp.; Witches Broom (WB), caused by *Moniliophthora perniciosa*, Cacao Swollen Shoot Virus (CSV), caused by a member of the genus Badnavirus; and Frosty Pod Rot (FPR), caused by *Moniliophthora roreri*. Some of the causal agents are globally distributed, but others have geographically restricted distribution (Gutiérrez et al., 2016; Marelli et al., 2019).

In Central America, more than half of the cacao production currently occurs in isolated rural areas on small-scale subsistence farms of fewer than 5 hectares. Consequently, the crop is seriously affected by the impact of diseases and the low-yielding potential of most plantations due to self and cross-incompatibility issues, pests, and diseases, as well as agronomical management. FPR and BPR are the two major diseases affecting cacao production, causing 30-100% yield losses (Phillips-Mora et al., 2006; Thevenin et al., 2012).

FPR disease was first officially reported in 1917 in Ecuador (Rorer, 1918); the fungus was formally named when this specimen was sent to R. Ciferri who ‘confirmed’ it as a new species of Monilia: *Monilia roreri* Cif. (Ciferri and Parodi, 1933). The disease is present in 13 countries in Latin America, including all countries of Central America (Sánchez-Mora et al., 2015). *M. roreri* only affects pods, with young pods with 2 to 3 months old being the most susceptible and dependent on climatic conditions (Sánchez and González, 1989). Farmers recognize *M. roreri* primarily by external symptoms on the fruits of cacao, especially by the appearance of signs of the pathogen, such as white mycelium, ashen in appearance, or the complete sporulation of the pathogen on the affected fruit tissue (Fig. 1A) (Phillips-Mora and Wilkinson, 2007). Although the origin of the pathogen remains unknown, recent findings with the help of molecular tools confirm that it was initially introduced in the coastal zone of Ecuador and the Magdalena Valley region in Colombia, which were areas of intensive production of the crop (Díaz-Valderrama et al., 2022).

**Fig. 1.**
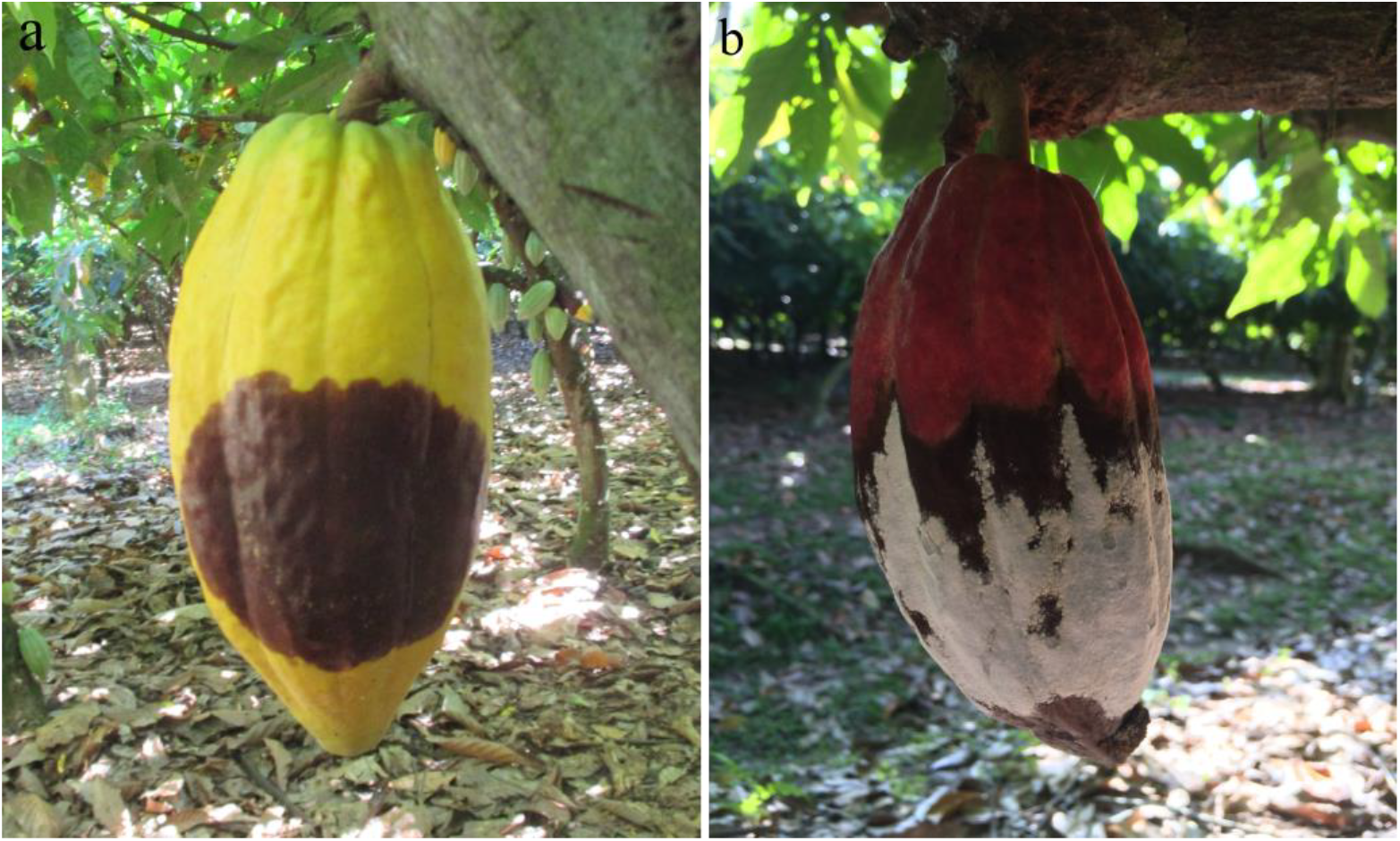
*Phytophthora* (a) and *Monilia* (b) disease in cacao pods.

**Fig. 2.**
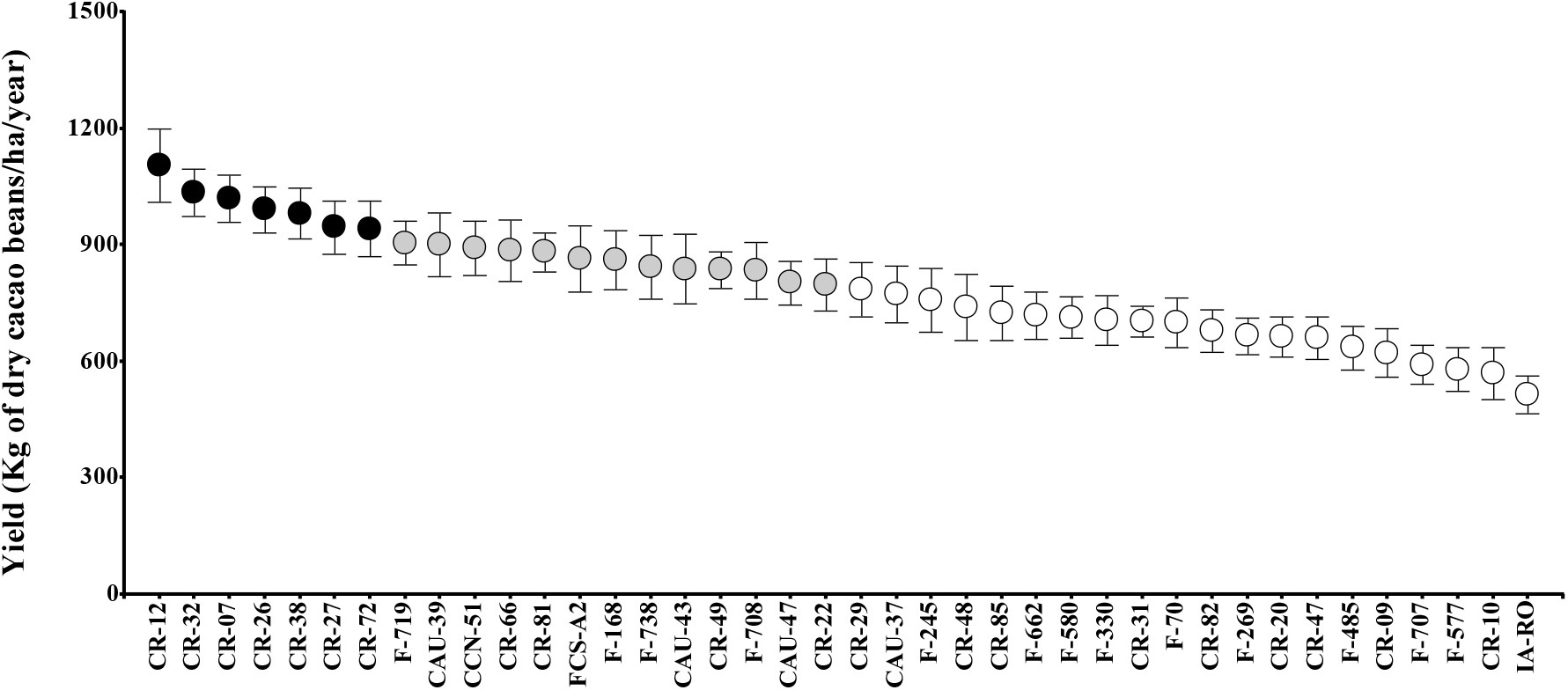
The average yield of dry cacao beans in kilograms per hectare (2013-20 17) of 40 cacao clones according to the Scott-Knott cluster test (p<0.05), different colors (black, gray, and white) represent statistical differences between groups. Data are means ± 95 % Standard Error of the mean

In the Central American region, *M. roreri* was first reported in Panamá in 1956, and after that, it was successively detected in Costa Rica in 1978, Nicaragua in 1980, Honduras in 1997, Guatemala in 2002, Belize in 2004, and Mexico in 2005 (Phillips-Mora et al. 2006; Phillips-Mora et al., 2012). As the disease was reported in each country, cacao production decreased considerably. For example, the total cacao production in Honduras in 1997 was approximately 5,500 tons; five years later, *M. roreri* arrived, the total cacao production decreased to 2,200 tons until reduced to 1000 tons in 2011 (FHIA, 2012). The impact on cacao production in the other Central American countries was like Honduras, especially because cacao producers were unaware of disease management and the genetic material used was not resistant to the disease.

Four species of Phytophthora cause BPR disease (*P*.*palmivora, P. megakarya, P. capcisi, and P. citrophtora*) *P. palmivora* being the most common (Drenth & Guest, 2004). *P. palmivora* is present in all cacao-producing areas (Ndubuaku and Asogwa, 2006). In Central America, BPR is caused mainly by *Phytophthora palmivora* (Ploetz, 2007); the disease spreads rapidly, covering the entire pod surface two weeks after infection. The disease mainly affects pods (Fig. 1B) but can also be observed in any part of the cacao plants. BPR is visually described as small, hard, dark lesions (Philip-Mora and Cerda, 2009). According to Marelli et al. (2019), BPR is responsible for losses of 873,000 tons of cacao per year worldwide, being the most harmful disease compared to other diseases affecting cacao production. Wet conditions like rainfall seasons, high relative humidity, and low temperature are the ideal allies of the disease (Dakwa, 1973)BPR is present in all Central American countries, and the incidence is higher when the fruit development stage coincides with ideal humidity and temperature conditions for the disease’s development.

One of the most critical challenges for a cacao breeding program is the evaluation time to select the desired traits in a new cacao clone. Generally, this evaluation is focused on yield and disease resistance and can take more than 10 years (Phillips-Mora et al., 2012). The selection of cacao clones is made by measuring the percentage of diseased fruits due to the natural incidence of the disease. However, this method does not discriminate between the clones with low or high production, contrasting with the selection of cacao clones by measuring the production potential. Furthermore, the potential production method does not consider the disease management cost, which is critical when analyzing economic profitability. A third selection method could be using a subjective index that combines yield and disease resistance, identifying cacao clones with high productivity and low disease incidence (Jaimez et al., 2020).

The Cacao breeding programs of Centro Agronómico Tropical de Investigación y Enzeñanza (CATIE) in Costa Rica and Fundación Hondureña de Investigación Agrícola (FHIA) in Honduras have developed new cacao clones with high resistance to FPR and BPR. (López et al., 2017; Phillips-Mora, 2015). These programs have validated the high yield and disease resistance using artificial methods. Posteriorly, clones were evaluated for disease incidence and severity under field conditions with the natural pressure of the inoculum.

This study aimed to compare three methods using Yield, Percentage Disease Pod (PDP), and Disease Production Index (DPI) as criteria to select cacao clones with high potential for production and tolerance to FPR and BPR, evaluating 40 cacao clones, 20 from CATIE and 20 from FHIA. Cacao clones evaluated were part of the project (“Programa Cacao Centro América” (PCC) developed between 2009-2015), selected by CATIE and FHIA breeding program for the cacao growers in Central America.

## 2. MATERIAL AND METHODS

### 2.1 Site, experimental design, and germplasm

The clones were planted in sandy, loamy soil (pH = 5.3) with low fertility (2.67 % of organic matter) and high iron content at the Experimental and Demonstration Center-Jesus Alfonso Sanchez (CEDEC-JAS) in La Masica, department of Atlántida (15°38’42.84” N, 87° 6’0.46” W, 25 m.a.s.l.) in the north of Honduras from 2013 to 2017. Weather conditions from 1986 to 2019 were reported as the annual mean temperature of 25.6 °C and 2,938.1 mm annual rainfall (Díaz et al., 2020).

One-year-old grafted cacao clones were planted in a square system of 3.5 m x 3.5 m, arranged in a complete randomized block design with four replicates and six plants per replicate. They were planted in agroforestry systems in association with different tropical wood species, mostly *Swietenia macrophylla, Cordia megaliths, Terminalia superba, Tabebuia rosea, Guarea grandifolia*, and *Ilex tectonic*, as permanent shade trees with no irrigation.

Mineral fertilizer was applied yearly with 136 kg of N-P-K (15-15-15), 45.4 kg of ammonium nitrate, and 45.4 kg of potassium chloride and lime amendments at a 0.5 tons/ha dose. Weather data on temperature, humidity, and precipitation were collected daily and reported as a monthly average.

Out of the forty clones used, twenty clones are part of FHIA’s Cacao and Agroforestry Program, and twenty clones are from the Cacao Breeding Program of CATIE (Table 1). At the FHIA cacao and agroforestry program, each cacao clone was evaluated for yield and disease incidence (FPR and BPR).

**Table 1.**
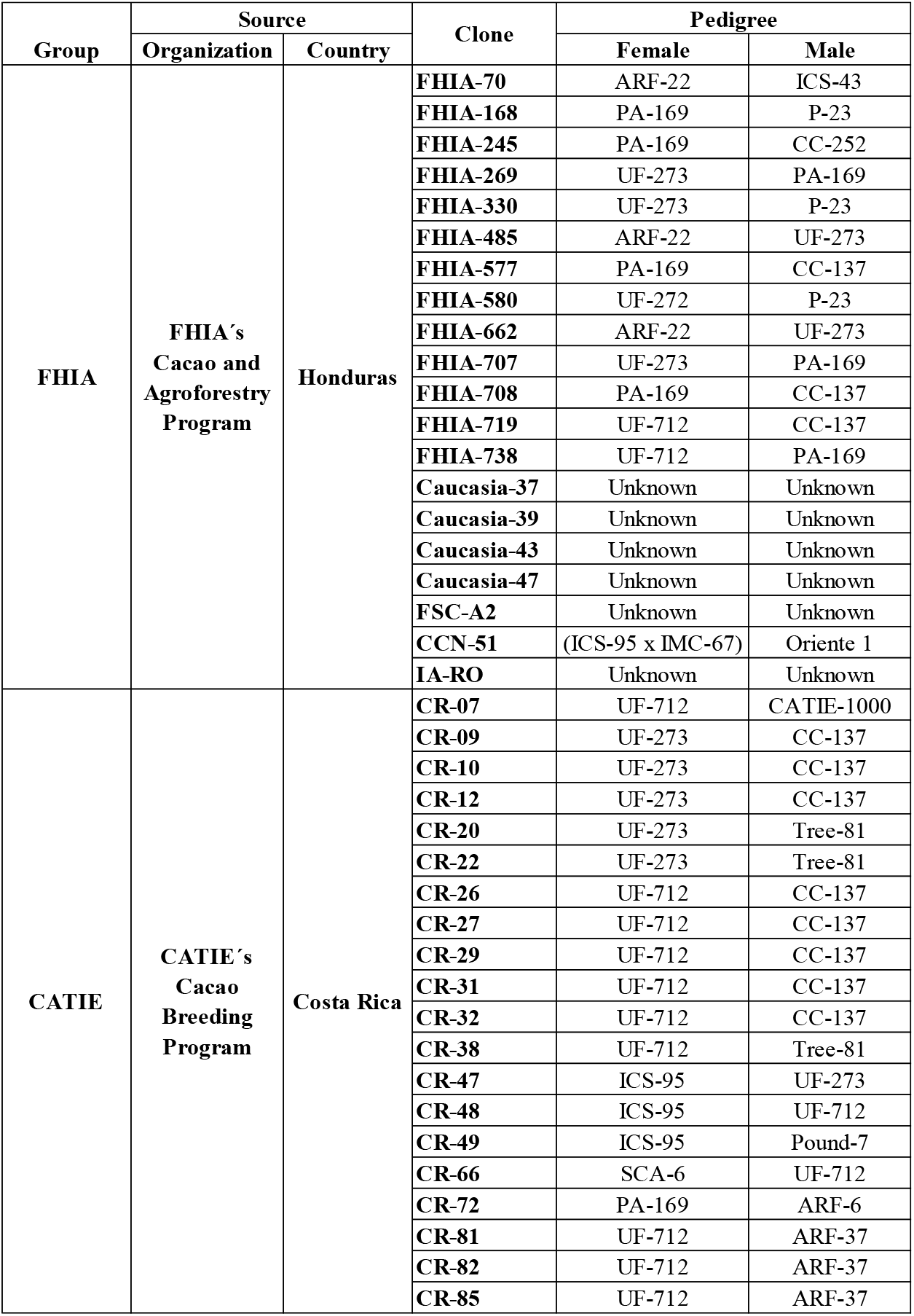
Cacao clones evaluated from CATIE and FHIA.

### 2.2 Variables evaluated

#### 2.2.1 Yield Variables

Yield was recorded as kg/ha of dry beans, also measured by the Pod Index (number of pods for 1 kg of the dry cacao bean) (IPGRI, 2000) and the Bean Index (Average weight of 100 dry cacao beans) (IPGRI, 2000).

#### 2.2.2 Diseases Variables

The Percentage of Disease Pod (PDP) was calculated as PDP = [NDP/ (NHP+NDP)] ×100, where NDP = annual number of diseased pods (calculated for Black Pod Rot, Frosty Pod Rot, and the two diseases together), NHP = annual Number of Healthy Pods (Jaimez et al., 2020).

#### 2.2.3 Index Variables

The Disease and Production Index (DPI) was calculated as follows: DPI= [(NHP+NDP) /DPC] ×0.1, where NHP = annual number of healthy pods, NDP = annual number of diseased pods (calculated separately for Black Pod Rot, Frosty Pod Rot, and two diseases together), and DPC = diseased pods coefficient. The DPC was calculated using the formula DPC = (NDP+1) /(NHP+1) (Jaimez et al., 2020). The DPI considers the effect of FPR, BPR, and FPR + BPR separately. A high DPI value is associated with the best cacao clones.

### 2.3 Statistical analysis

Data analysis was performed using the InfoStat software (Di Rien zo et al., 2020). The statistical difference was determined using the one-way ANOVA method followed by the Scott-Knott test method for grouping means of cacao clones, as a large number of treatments were analyzed (Jaimez et al., 2020). The results were expressed as the mean ± Standard Error (SE). In addition, Spearman correlation and Principal Components Analysis (PCA) were carried out for yield, PDP, and DPI variables using R statistical software (R core team, 2019) through the Corrplot package (Wei et al., 2017). For correlation analysis, Factoxtra (Kassambra and Mundt, 2020) and ggplot2 (Vu, 2020) for biplot of principal component analysis.

## 3. RESULTS

### 3.1 Yield

All cacao clones evaluated individually showed yields above 500 kg/ha. The Scott-Knott clusterization analysis shows that the yield variable formed three different groups (Fig.). Seven cacao clones showed to be the most productive (CR-12, CR-32, CR-07, CR-26, CR-38, CR-27, and CR-72), thirteen cacao clones were clustered in the second group, and twenty cacao clones in the last group. Boxplot showed that the yield average of all cacao clones was very similar throughout the 2013-2017 period (Fig. 3**A**), showing a decreased yield in 2014; furthermore, the results showed that Bean and Pod Indexes were negatively correlated; when the Bean Index increases, the Pod Index decreases (Fig. 3**B**). The correlation between Bean and Pod Indexes is essential for the final selection of cacao clones since it is directly associated with the dry cacao bean yield per hectare.

**Fig. 3.**
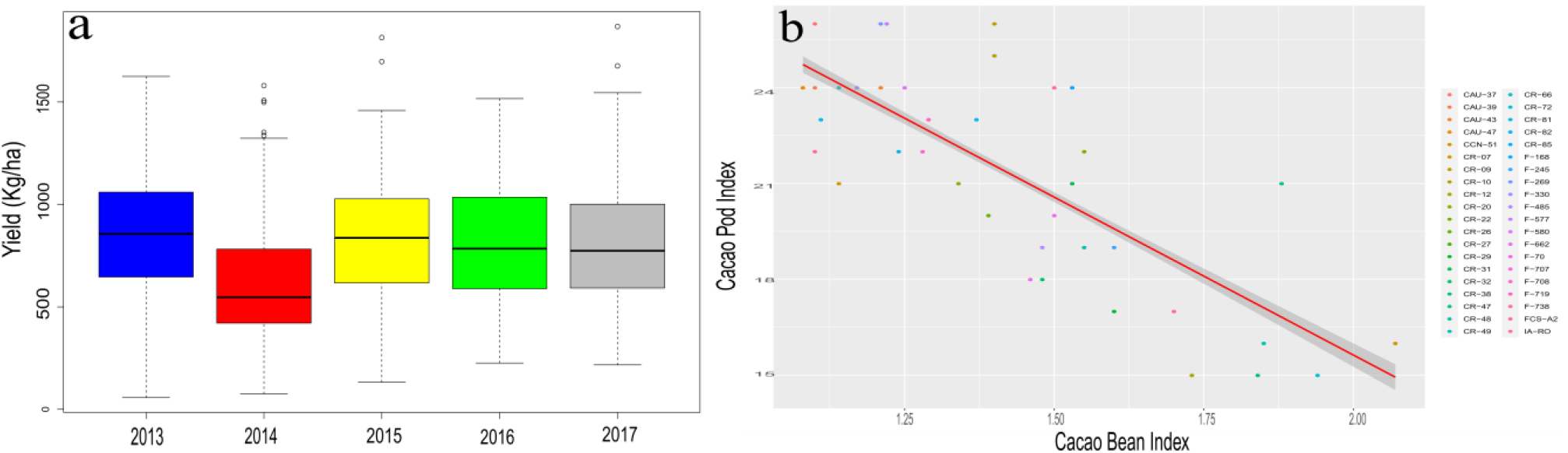
Yield components. Boxplot representing total cacao yield per year (A), the relation between Pod Index and Bean Index (B)

### 3.2 Percentage Disease Pod (PDP)

The incidence of BPR and FPR observed was dissimilar in the evaluation period. In general, there was a higher incidence of BPR than FPR. The number of diseased pods with BPR increased from 2013 to 2017 (Fig. 4**A**). The over-the-year averages of BPR ranked from 20 to 55 % of disease incidence. Therefore, the clones could be statistically separated into two groups (Fig. 5). The first group has a range of Percentage Disease Pods between 33.34-52.94 % (Black points), and the second group has a range of Percentage Disease Pods between 16.64-31.22 % (White points). The incidence of FPR remained steady throughout the years (Figure 4B), maintaining the incidence of FPR disease lower than 5 % during all years without differences between cacao clones (Fig. 6). Thus, we see a marked difference in the Percentage of Disease Pods. Furthermore, more incidence of BPR was observed when environmental conditions such as rain and temperature were high in the last quarter of the year, which is typical in tropical regions.

**Fig. 4.**
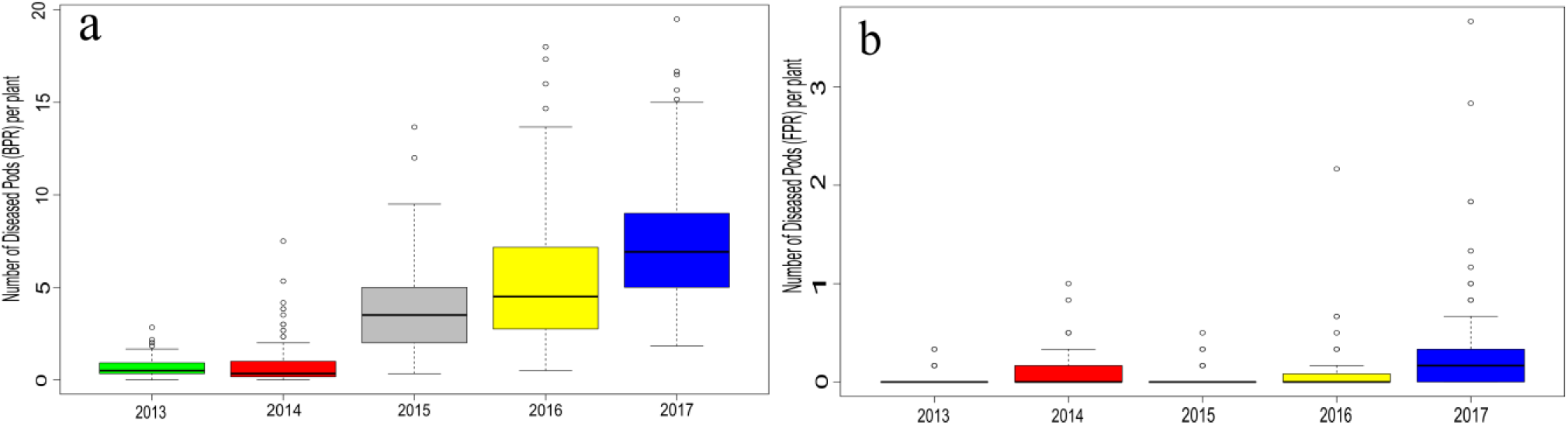
Diseased pods per plant between 2013 and 2017. Black Pod Rot (BPR) (A), and Frosty Pod Rot (FPR) (B)

**Fig. 5.**
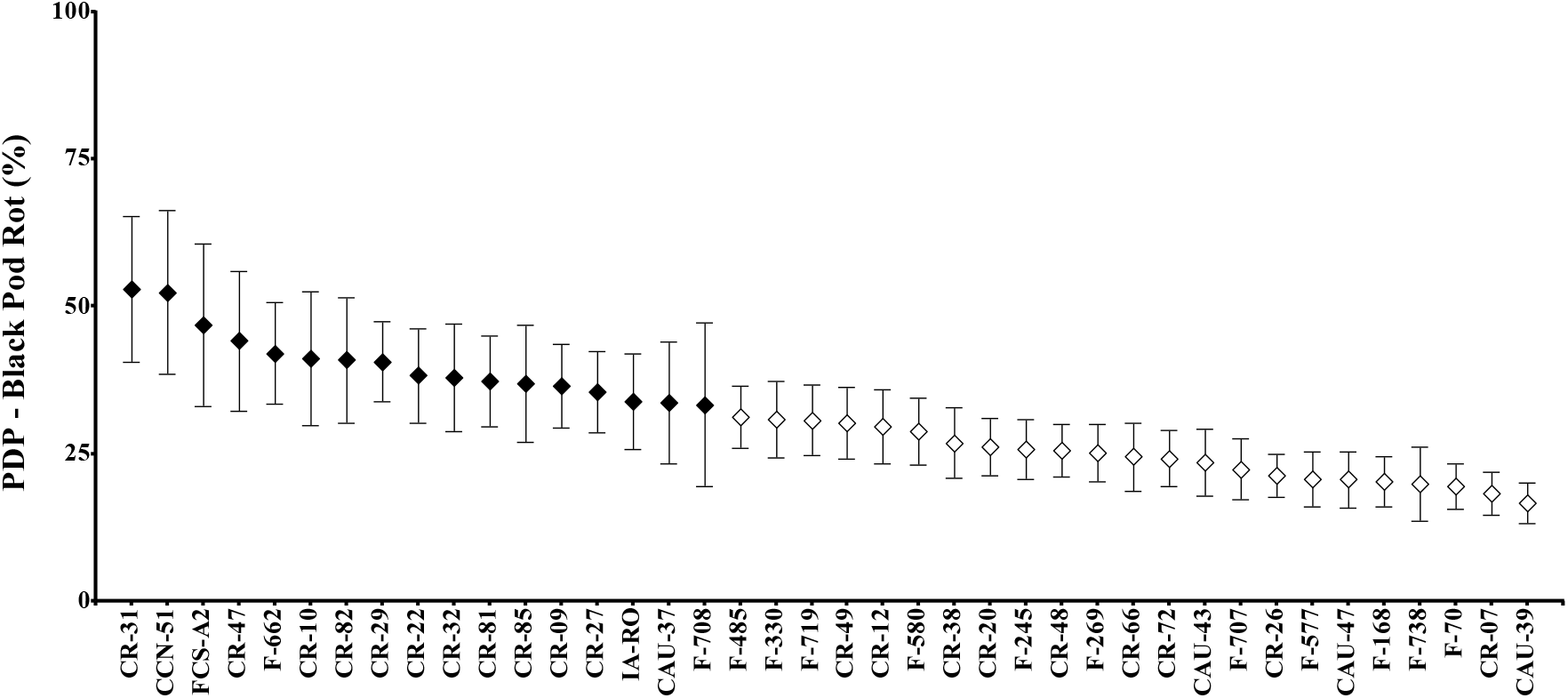
The percentage of pods affected by BPR, different colors (white and black) in the boxplot shows statistical differences according to the Scott-Knott test (p<0.05). Data are means ± 95 % Standard Error of the mean

**Fig. 6.**
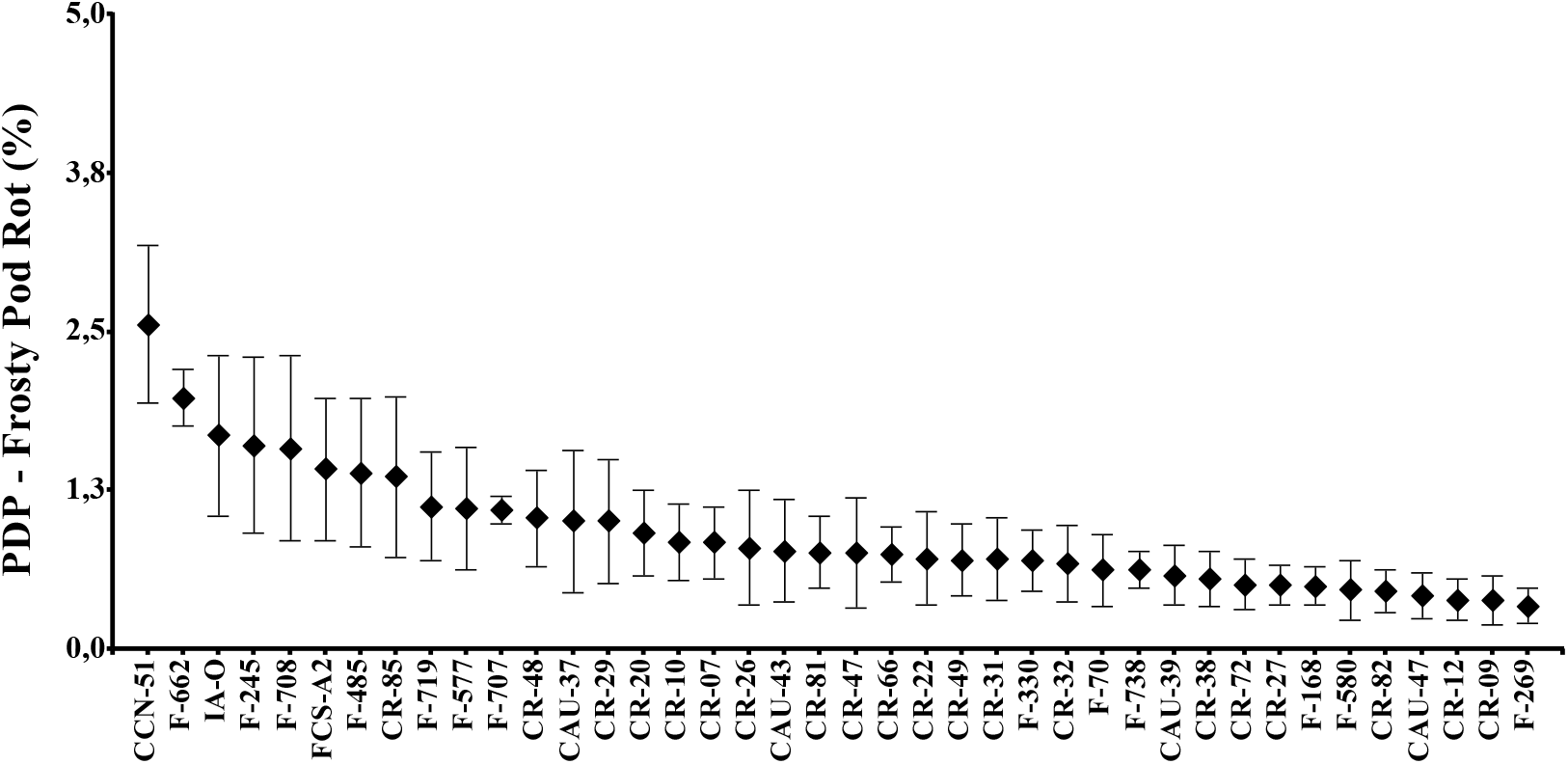
According to the Scott-Knott test, the percentage of pods affected by FPR does not show statistical differences between cacao clones (p<0.05). Data are means ± 95 % Standard Error of the mean.

### 3.3 Disease and Production Index (DPI)

The Scott-Knott test showed that the DPI for FPR was divided into two groups (p <0.05); in the first group, 20 cacao clones were observed, which were less affected by FPR, whereas in the second group, the cacao clones with more incidence of the disease (Fig. 7A).

**Fig. 7.**
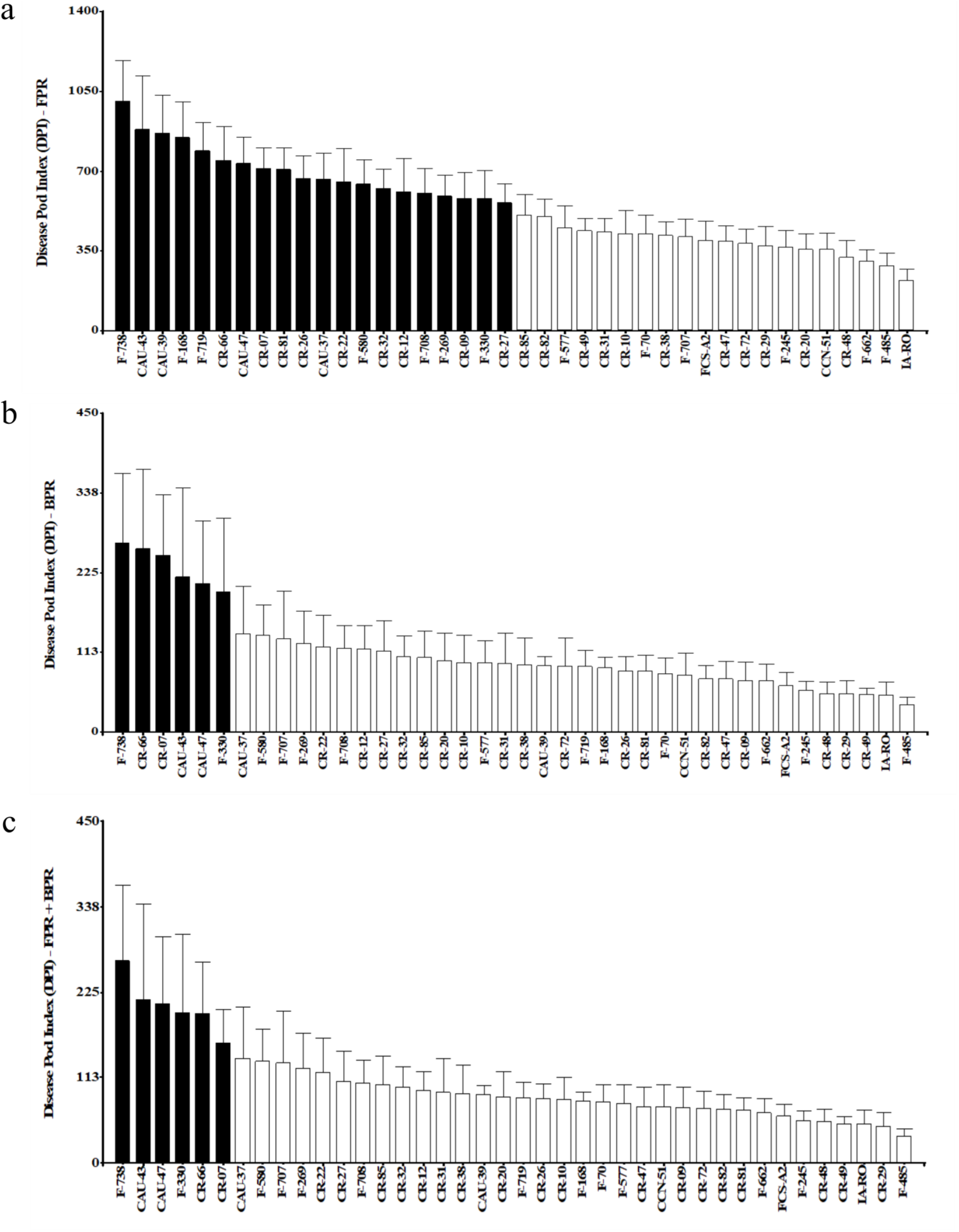
Disease and Production Index Pod (DPI). A) FPR, B) BPR, and C). FPR + BPR. Different colors (black and white) represent statistical differences between groups (p> 0.05) according to the Scott-Knott test.

The DPI for BPR was divided into two groups (p <0.05). The first group of cacao clones, F-738, CR-66, CR-07, CAU-43, CAU-47, and F-330, distinguished among the cacao group clones as the lowest incidence of BPR (Fig. 7B). When the DPI was calculated considering the incidence of the two diseases (FPR and BPR), the cacao clones aggregated statistically in two groups(p<0.05). We observed the highest DPI values in the clones F-738, CAU-43, CAU-47, F-330 CR-66, and CR-07 (Fig. 7C), compared with the rest of the cultivars.

### 3.4 Correlation and PCA analysis

Correlation and PCA analysis were performed between production variables (Pod index, Bean Index, and Yield), disease variables (Percentage Disease Pod (PDP), Total Disease Pod (TDP), Total Healthy Pod (THP)), and Disease and Production Index variables (DPI-FPR, DPI-BPR, and DPI(FPR+BPR)).

Figure 8A shows different results of the correlation analysis. For the Yield variables, a positive correlation between Yield and Total Healthy Pod (THP) (R=0.92) is present, and there is a high negative correlation between the seed index and the pod index (R = -0.71). In the disease variables, various positive and negative correlations were found. As expected, Total number of Healthy Pods (THP) correlates positively with yield (R = 0.92). It also correlates positively with disease and production index (DPI- (FPR + BPR) (R = 0.51), DPI-FPR (R = 0.85), and DPI-BPR (R = 0.50). The variable of the total number of diseased pods (TDP) correlates positively with the variable of the percentage of diseased pods (TDP) (R = 0.94). However, a negative correlation is observed with the variables of Total Disease Pod (TDP) with DPI-(FPR + BPR) (R = -0.69) and with DPI-BPR (R = -0.7). The yield variable positively correlates with DPI-FPR (R = 0.78), possibly due to the low incidence of Frosty Pod Rot disease observed during the investigation. The variable Percentage of Disease Pods (PDP) showed a high negative correlation with the variables DPI (FPR + BPR) (R = -0.88) and DPI-BPR (R = -0.89). Finally, a positive correlation between DPI- (FPR + BPR) and DPI-BPR (R = 1) is shown. Using the PCA analysis, the first two dimensions explained 62.65% of the overall variation (Fig. 8B). The variation related to the first component was primarily associated with yield (THP, Yield, and DPI-FPR) and DPI (DPI-BPR and DPI-(FPR+BPR)) variables. The second component was associated with disease variables (PDP and TDP). It was observed that the DPI variables are more associated with Disease than yield variables.

**Fig. 8.**
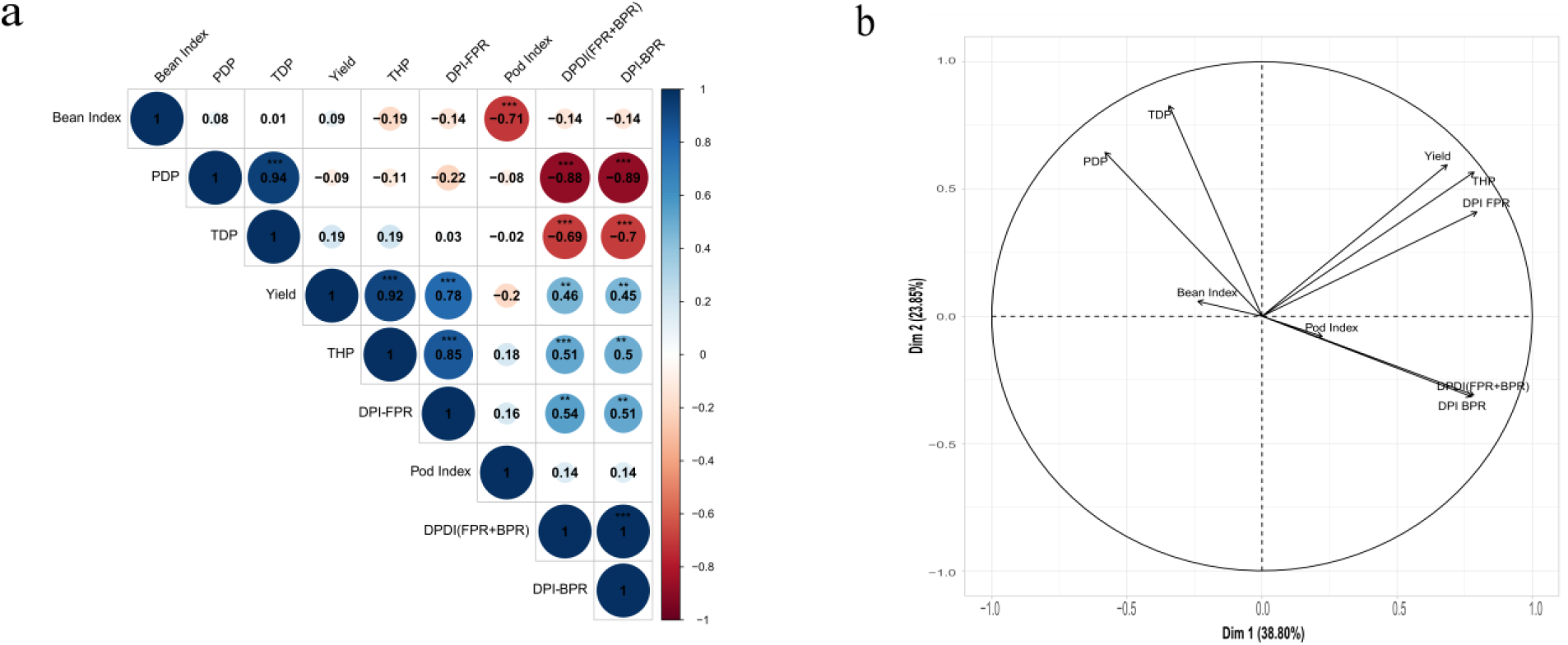
Relation between Disease (PDP, TDP, and THP), Yield (Yield, Bean Index, and Pod Index), and DPI (DPI-FPR, DPI-BPR, and DPI-(FPR+BPR) variables. Correlation Analysis (A), Principal Components Analysis (B)

## 4. DISCUSSION

### 4.1 Genetic improvement for cacao clones with high yield and disease-resistant

Handing a new cacao clone to growers with high productivity and resistance to pests and diseases is the dream of a cacao breeder. This task is never easy to achieve due to the lengthy selection process and time that this activity entails. Pests and diseases can destroy 20–30% or more of total cacao production, and high yield and disease resistance have received the most attention from breeders (Gutiérrez et al. 2016; Lopes et al. 2011). The primary strategy of the cacao genetic improvement programs is based on the recurrent selection using parental trees with high yield and resistance to pests and diseases. Those recurrent selections in cacao breeding programs are continuing with a broader level of diversity and aiming at accumulating favorable alleles for yield and resistance to the four diseases causing the most damage in cacao plantations, moniliasis (*Moniliophthora roreri*), Witches broom (*Moniliophthora perniciosa)*, black pod disease (*Phytophthora* sp.) and mal del machete (*Ceratocystis* sp.) (Baudouin 1997; Rodriguez-Medina et al., 2019; Lopes et al., 2011; Gutierrez et al., 2016). Obtaining new cacao clones resistant to these diseases with desirable agronomic characteristics, as well as sensory qualities that contribute to obtaining adequate chocolate quality and more income for cacao producers, is the main objective of genetic improvement at present (Pimenta Neto et al., 2018). The choice of selecting desirable parents for cacao breeding traditionally depended on the availability and characterization of germplasm. However, polygenic traits significantly influenced by the environment are more challenging to measure without a tool to identify the significant genes influencing the phenotype (Bekele and Phillips-Mora, 2019). Nowadays, with molecular biology tools, it is possible to identify molecular markers associated with resistance genes FPR and BPR and the characterization of germplasm banks that could be used for genetic improvement in the future (Osorio-Guarin et al., 2020; Rodriguez-Polanco et al., 2020; Legavre et al., 2015).

The two old and most extensive collections of cacao germplasms are in Trinidad and Tobago (CRU/UWI) and Costa Rica at CATIE, and both are used as a source of germplasm for genetic improvement (Monteiro et al., 2009; Laliberté et al., 2012). At the same time, other more recent germplasm collections, such as FHIA in Honduras, have a core of genetic material as a source of specific characteristics of resistance to diseases and yield (Somarriba and Villalobos, 2013; Somarriba et al., 2013). Therefore, 40 cacao clones from the two genetic improvement programs (FHIA and CATIE) were evaluated in the present study. This study aimed to select the best clones with resistance to BPR, FPR, and high productivity as an initiative to transfer the best clones to Central American growers.

The clones evaluated showed more resistance to FPR than to BPR based on natural incidence. The reason may be that when FPR arrived in Central America about 20 years ago, its appearance was so devastating that genetic improvement programs focused their efforts on selecting cacao clones with resistance to FPR, and genetic improvement for resistance to BPR was neglected (FHIA 2012; Phillips-Mora et al. 2005, 2006, 2012).

In contrast, while resistance to BPR has been identified in several germplasm accessions, resistance to FPR is relatively uncommon. In an evaluation of seventy new cacao clones, Phillips-Mora and Castillo (1999) report that only two clones (3%) were characterized as moderately resistant (MR). Furthermore, based on their screening results, only ten (2.3%) out of 441 clones (56%) of the CATIE collection were identified as resistant or moderately resistant to FPR. Phillips-Mora et al. (2017) stated that out of 1260 clones from the CATIE collection, 76 (6%) showed tolerance to FPR. On the other hand, using molecular tools, Gutierrez et al., 2021 found and reconfirmed QTLs associated with FPR and BPR resistance and the expression of genes related to plant defense and disease resistance.

There are several challenges for cacao breeders, among them low productivity, the higher pressure of pests and disease due to climate change, small production units, high production costs, and above all, maintaining the quality of the final product so that consumers can be satisfied (Bekele and Phillips-Mora, 2019). Cacao breeding programs should design strategies that include all the possible variables, considering the long process of improving a new cacao cultivar. The main objective of a development program is to create new cacao cultivars with high yield and disease resistance. However, there are subdivisions within each of these desirable features, as they all come together to achieve the objective. For example, among the yield components, a new cultivar should have cacao fermented seeds > 1g, a low pod index, a high number of seeds per pod, and high content of butterfat among others (Soria, 1977). Other selection criteria include vigor, self-compatibility, uniform plant type, compactness in tree size, precocity (early flowering and maturing) tolerance to drought stress, and quality expressed in terms of bean flavor, purity, and food safety. (Ahnert 2009; Bekele and Phillips-Mora, 2019). The second important objective is genetic disease resistance because this represents the most serious biological constraint in cacao production (Gutierrez et al., 2016).

### 4.2 Relation between Yield, Percentage Disease Pod, and Disease Pod Index

Cacao production and its components should be appropriately appraised since they are considered polygenic characters and, therefore, highly influenced by environmental factors (Monteiro et al., 2009). Consequently, yield continues to be the main objective in a cacao production unit; as a result, the growers use all the technologies available to achieve higher yields, including germplasm, fertile soils, planting distance, fertilization, irrigation, and self-compatibility to make the crop profitable. However, yield can be reduced without effective pest and disease control measures. In our study, the statistical analysis was carried out for the variable yield of seven cacao clones (CR-12, CR-32, CR-07, CR-26, CR-38, CR-27, and CR-72) (Fig.). The results showed how the environmental factor is significantly driving the yield. Cacao clones evaluated in Costa Rica by CATIE showed higher yields than those evaluated in Honduras by FHIA (Arciniegas et al., 2005). The high yield is a characteristic that could be inherited from the parent of the clone (CC-137 characterized as a high yield, low incidence of diseases, low index pod, and long grain) since five of these clones have the clone CC-137 as a mother parent (Arciniegas et al., 2005). In our study, the yield that CR-12, CR-32, CR-07, CR-26, CR-38, CR-27, and CR-72 showed (950-1150 kg/ha) is acceptable considering that the global average is between 300-400 kg/ha. Similar results were found in other studies evaluating elite cacao clones under field disease pressure for at least four years (Ofori et al., 2019; Sánchez-Mora et al., 2015; Solis et al., 2015).

Another way to quantify cacao production is to consider the pod number affected by diseases and their effect on the total production. In our study, all the cacao clones used showed a high level of genetic resistance to FPR, and there was no difference between the clones (Fig.); however, a high percentage of pods were affected by BPR (Fig.) and within the group of clones evaluated, two groups with different infection levels were observed. Since BPR is a disease in all cacao-producing areas of the world (Ploetz, 2007), it is unsurprising that BPR causes more damage in some humid seasons than FPR (Phillips-Mora and Cerda, 2009). However, in this case, the clones have a higher genetic resistance to FPR than BPR.

Since chemical control is not commonly used in managing FPR and BPR, selecting genetically resistant cacao clones is the most effective strategy. It also helps to avoid environmental contamination by reducing the use of pesticides. In this sense, it is still a global challenge to develop cacao clones with resistance to different BPR strains (Purwantara et al., 2015; Mcmahon et al., 2015; Fister et al., 2020; Decloquement et al., 2021) although, more resistant clones to FPR have been found (Phillips-Mora et al., 2012; Torres et al., 2011).

Yield and Percentage of Disease Pods can be used as criteria for cacao clone selection; however, the percentage of diseased pods has limitations because it does not discriminate between genotypes with high and low production potential (Jaimez et al., 2020). In this sense, it can be most useful as a method that includes both criteria for yield and percentage of the diseased pod as an index for cacao clone selection. Some studies have been carried out to select new cacao clones using both criteria: yield and percentage of diseased pods (Nyassé et al., 2003; Ofori et al., 2020; Cervantes-Martinez et al., 2006; Efombagn et al., 2007, 2011). On the other hand, this criteria combination has been used in other species such as *Zea mays* (Horne et al., 2016), *Saccharum officinarum* (Magarey et al., 2003a), *Arachis hypogaea* L. (Iroume and Knauft 1987), *Cicer arietinum* (Toker and Çanci 2003), *Capsicum annuum* L. (Sreenivas et al., 2020).

Jaimez et al, (2020) developed a disease and production index (DPI) for the selection of cacao clones highly productive and tolerant to pod rot diseases. When we evaluated this index for cacao clone classification, two groups were formed especially when Yield, BPR, and FPR were included (Fig.). This index is valid for breeders and growers because it balances cacao crops’ production potential and disease resistance. The cacao clones that combine the best yield and disease resistance have the highest index value.

### 4.3 Correlation analysis, PCA, and final selection

Finally, a correlation analysis between production potential and disease variables was carried out (Fig.). The analysis results are consistent with those shown in selecting cacao clones by yield, Percentage of Disease Pod, and Disease Pod Index (DPI).

As shown in the PCA analysis, the pod index variable negatively correlated with the seed index, an important characteristic to use as a criterion for selecting new cacao clones. The variables yield, Total Healthy Pod (THP), and Total Diseased Pods (TDP) were grouped as the main factors contributing to cacao yield performance. In this group, the Disease and Production Index for Frosty Pod Rot (DPI-FPR) correlated due to the low incidence of FPR in this study. On the other hand, DPI-BPR and DPI-(FPR+BPR) are on the opposite side of PDP and TDP, suggesting that both indexes depend on PDP and TDP.

In the correlation analysis, variables for cacao yield performance correlated positively, and index variables correlated negatively with disease variables. Both analyses were complemented in their results (Fig.). The final decision in the cacao clone’s selection process must always be balanced, including yield and disease resistance components. The cacao clones that show the highest yield are not necessarily the most resistant to diseases because the genetic yield potential (Ofori et al., 2020) is different from the accumulation of genes with the total or partial expression of resistance to pests and diseases and have environmental influence (Nyadanu et al., 2017).

Finally, after a lengthy selection period, the breeder should select the best cacao clones according to the objectives of his genetic improvement program. Table 2 presents the top 10 best cacao clones, selected using the three criteria (Yield, Diseases, and disease and production index (DPI)) as an example of how to make the final selection. Note that in the disease and DPI criteria, there are subdivisions of different criteria according to the disease for which we are interested in directing our selection. In our study, cacao clones selected by yield criteria differ from those selected using the other two criteria because those cacao clones with high yield do not always associate with disease resistance. The final decision on which method to use depends on the breeding program objectives and the environmental conditions in which the cacao clones selected will be planted. Environmental conditions highly influence disease incidence. However, in tropical areas, production coinciding with high humidity and low temperatures suggests using a Disease Percentage or Disease and Production Index (DPI) criteria.

**Table 2.** The top 10 cacaos were selected using three criteria (Yield, Disease Percentage, and Disease and Production Index (DPI))

| No | YIELD | DISEASE PERCENTAGE |  |  | DISEASE AND PRODUCTION INDEX (DPI) |  |  |
| --- | --- | --- | --- | --- | --- | --- | --- |
|  |  | FPR | BPR | FPFR+BPR | DPI-FPR | DPI-BPR | DPI-(FPR+BPR) |
| 1 | CR-12 | F-269 | CAU-39 | CAU-39 | F-738 | F-738 | F-738 |
| 2 | CR-32 | CR-09 | CR-07 | F-738 | CAU-43 | CR-66 | CAU-43 |
| 3 | CR-07 | CR-12 | F-70 | CR-07 | CAU-39 | CR-07 | CAU-47 |
| 4 | CR-26 | CAU-47 | F-738 | F-70 | F-168 | CAU-43 | F-330 |
| 5 | CR-38 | CR-82 | F-168 | CAU-47 | F-719 | CAU-47 | CR-66 |
| 6 | CR-27 | F-580 | CAU-47 | F-168 | CR-66 | F-330 | CR-07 |
| 7 | CR-72 | F-168 | F-577 | F-577 | CAU-47 | CAU-37 | CAU-37 |
| 8 | F-719 | CR-27 | CR-26 | F-707 | CR-07 | F-580 | F-580 |
| 9 | CAU-39 | CR-72 | F-707 | CAU-43 | CR-81 | F-707 | F-707 |
| 10 | CCN-51 | CR-38 | CAU-43 | CR-26 | CR-26 | F-269 | F-269 |

## 5. CONCLUSION

The selection of new cacao clones with high yield and resistance to diseases is not an easy task because environmental conditions and time influence this process. In our study, we showed that it is possible to make the final selection of cacao clones using three selection criteria according to the objectives of the breeding program. Although selection using the criteria for yield or disease incidence is the most common, it is also possible to use a Disease and Production Index (DPI) that combines the two previous criteria for the final decision to select new cacao clones for growers.

## Authorship contribution statement

**Marlon Enrique López:** Conceptualization, formal analysis, data curation, writing, original draft (Review and editing), **Oscar Arnulfo Argueta**: Investigation, methodology, formal analysis, data curation, **Aroldo Dubón**: Conceptualization, Investigation, methodology, field data, resources, **Francisco Javier Díaz**: Funding acquisition, supervision, writing (Editing), resources.

## Acknowledgments

The author thanks the financial support of the “Programa Cacao Centro América” (PCC) project and acknowledges the scientific staff of FHIA and CATIE for helping to develop this project.

## REFERENCES

Arciniegas Leal, A. M. (2005). Caracterización de árboles superiores de cacao (Theobroma cacao L.) seleccionados por el programa de mejoramiento genético del CATIE.

Ahnert, D. (2009). Ideotype breeding in cocoa. In Proceedings of the international workshop on cocoa breeding for farmers’ needs (pp. 157–166).

Baudouin, L., Baril, C., Clément-Demange, A., Leroy, T., & Paulin, D. (1997). Recurrent selection of tropical tree crops. Euphytica, 96(1), 101–114.

Bekele, F., & Phillips-Mora, W. (2019). Cacao (Theobroma cacao L.) breeding. In Advances in plant breeding strategies: Industrial and food crops (pp. 409–487). Springer, Cham.

Ciferri, R. and Parodi, E. (1933) Descrizione del fungo che causa la “Moniliasis” delcacao.Phytopath. Z.6, 539– 542.

Cervantes-Martinez, C., Brown, J. S., Schnell, R. J., Phillips-Mora, W., Takrama, J. F., & Motamayor, J. C. (2006). Combining ability for disease resistance, yield, and horticultural traits of cacao (Theobroma cacao L.) clones. Journal of the American Society for Horticultural Science, 131(2), 231–241.

Díaz-Valderrama, J. R., Zambrano, R., Cedeño-Amador, S., Córdova-Bermejo, U., Casas, G. G., García-Zurita, N., … & Aime, M. C. (2022). Diversity in the invasive cacao pathogen Moniliophthora roreri is shaped by agriculture. Plant Pathology, 71(8), 1721–1734.

Drenth, A., & Guest, D. I. (2004). Diversity and management of Phytoph vthora in Southeast Asia. Diversity and Management of Phytophthora in Southeast Asia.

Dakwa, J. T. (1973). The yield and disease patterns in a black pod resistance trial in Ghana. Ghana Journal of Agricultural Science, 6, 205–11.

Díaz, F.J., Dubon, Á., Martínez, A., 2020. Registros climaticos ’ del CEDEC-JAS y CADETH. P. 5-9. In: Informe Técnico 2019 Programa de Cacao y Agroforestería. Fundacion ’ Hondurenãde Investigacion Ágrícola. La Lima, Cortés, Honduras, C.A., p. 89.

Di Rienzo JA, Casanoves F, Balzarini MG, Gonzalez L, Tablada M, Robledo CW, 2020. InfoStat versión 2020p. Centro de Transferencia InfoStat, FCA, Universidad Nacional de Córdoba, Argentina.

Decloquement, J., Ramos-Sobrinho, R., Elias, S. G., Britto, D. S., Puig, A. S., Reis, A., … & Marelli, J. P. (2021). Phytophthora theobromicola sp. nov.: a new species causing black pod disease on cacao in Brazil. Frontiers in microbiology, 12.

Efombagn, M. I. B., Nyassé, S., Sounigo, O., Kolesnikova-Allen, M., & Eskes, A. B. (2007). Participatory cocoa (Theobroma cacao) selection in Cameroon: Phytophthora pod rot-resistant accessions identified in farmers’ fields. Crop Protection, 26(10), 1467–1473.

Efombagn, M. I. B., Bieysse, D., Nyassé, S., & Eskes, A. B. (2011). Selection for resistance to Phytophthora pod rot of cocoa (Theobroma cacao L.) in Cameroon: Repeatability and reliability of screening tests and field observations. Crop Protection, 30(2), 105–110.

Evans, H. C. (2007). Symposium Cacao Diseases: Important Threats to Chocolate Production Worldwide Cacao Diseases-The Trilogy Revisited. 10.1094/PHYTO-97-12-1640

Fister, A. S., Leandro-Muñoz, M. E., Zhang, D., Marden, J. H., Tiffin, P., dePamphilis, C., … & Guiltinan, M. J. (2020). Widely distributed variation in tolerance to Phytophthora palmivora in four genetic groups of cacao. Tree Genetics & Genomes, 16(1), 1–9.

FHIA. (2012). La Moniliasis de Cacao: El Enemigo A Vencer. La Lima, Cortés, Honduras, C.A.

Gutiérrez, O. A., Campbell, A. S., & Phillips-Mora, W. (2016). Breeding for disease resistance in cacao. In Cacao Diseases: A History of Old Enemies and New Encounters (pp. 567–609). Springer International Publishing. 10.1007/978-3-319-24789-2_18

Gutiérrez, O. A., Campbell, A. S., & Phillips-Mora, W. (2016). Breeding for disease resistance in cacao. In Cacao Diseases (pp. 567–609). Springer, Cham.

Gutiérrez, O. A., Puig, A. S., Phillips-Mora, W., Bailey, B. A., Ali, S. S., Mockaitis, K., … & Motamayor, J. C. (2021). SNP markers associated with resistance to frosty pod and black pod rot diseases in an F1 population of Theobroma cacao L. Tree Genetics & Genomes, 17(3), 1–19.

Horne, D. W., Eller, M. S., & Holland, J. B. (2016). Responses to recurrent index selection for reduced Fusarium ear rot and lodging and increased yield in maize. Crop Science, 56(1), 85–94.

Iroume, R. N., & Knauft, D. A. (1987). Heritabilities and correlations for pod yield and leafspot resistance in peanut (Arachis hypogaea L.): Implications for early generation selection. Peanut Science, 14(1), 46–50.

Jaimez, R. E., Vera, D. I., Mora, A., Loor, R. G., & Bailey, B. A. (2020). A disease and production index (DPI) for the selection of cacao (Theobroma cacao) clones are highly productive and tolerant to pod rot diseases. Plant Pathology, 69(4), 698–712. 10.1111/ppa.13156

Kassambra A, Mundt F (2020). Factoextra: extract and visualize the results of multivariate data analyses. https://cran.rproject.org/web/packages/factoextra/index.html.

López, M., Ramirez, O., & Dubón, A. (2017). Catálogo De Cultivares De Cacao (Theobroma cacao L.) Evaluados y Seleccionados por la FHIA.

Legavre, T., Ducamp, M., Sabau, X., Argout, X., Fouet, O., Dedieu, F., … & Lanaud, C. (2015). Identification of Theobroma cacao genes differentially expressed during Phytophthora megakarya infection. Physiological and Molecular Plant Pathology, 92, 1–13.

Laliberté, B., Cryer, N. C., Daymond, A. J., End, M. J., Engels, J. M., Eskes, A., … & Weise, S. (2012). A global strategy for the conservation and use of cacao genetic resources, as the foundation for a sustainable cocoa economy.

Lopes, U. V., Monteiro, W. R., Pires, J. L., Clement, D., Yamada, M. M., & Gramacho, K. P. (2011). Cacao breeding in Bahia, Brazil: strategies and results. Crop breeding and applied biotechnology, 11, 73–81.

Monteiro, W. R., Lopes, U. V., & Clement, D. (2009). Genetic improvement in cocoa. In Breeding plantation tree crops: tropical species (pp. 589–626). Springer, New York, NY.

Marelli, J. P., Guest, D. I., Bailey, B. A., Evans, H. C., Brown, J. K., Junaid, M., Barreto, R. W., Lisboa, D. O., & Puig, A. S. (2019). Chocolate is under threat from old and new cacao diseases. Phytopathology, 109(8), 1331–1343. 10.1094/PHYTO-12-18-0477-RVW

McMahon, P. J., Susilo, A. W., Parawansa, A. K., Bryceson, S. R., Mulia, S., Saftar, A., … & Keane, P. J. (2018). Testing local cacao selections in Sulawesi for resistance to vascular streak dieback. Crop Protection, 109, 24–32.

Magarey, R.C., Kuniata, L.S., Croft, B.J., Chandler, K.J., Irawan Kristini, A., Spall, V.E., Samson, P.R. and Allsopp, P.G. (2003a). International activities to minimise industry losses from exotic pests and diseases. Proc. Aust. Soc. Sugar Cane Technol., 25: (CDROM), 9 p.

Nyassé, S., Efombagn Mousseni, I. M.w, Bouambi, E., Ndoumbe-Nkeng, M., & Eskes, A. B. (2003). Early selection for resistance to Phytophthora megakarya in local and introduced cocoa varieties in Cameroon. Tropical Science, 43(2), 96–102.

Nyadanu, D., Akromah, R., Adomako, B., Akrofi, A. Y., Dzahini-Obiatey, H., Lowor, S. T., … & Attamah, P. (2017). Genetic control, combining ability and heritability of resistance to stem canker in cacao (Theobroma cacao L.). Euphytica, 213(12), 1–13.

Ndubuaku TCN, Asogwa EU (2006) Strategies for the control of pests and diseases for sustainable cocoa production in Nigeria. Afr Sci 7:202–216

Osorio-Guarín, J. A., Berdugo-Cely, J. A., Coronado-Silva, R. A., Baez, E., Jaimes, Y., & Yockteng, R. (2020). Genome-wide association study reveals novel candidate genes associated with productivity and disease resistance to Moniliophthora spp. in cacao (Theobroma cacao L.). G3: Genes, Genomes, Genetics, 10(5), 1713–1725.

Ofori, A., Arthur, A., & Padi, F. K. (2019). Extending the cacao (Theobroma cacao L.) gene pool with underrepresented genotypes: growth and yield traits. Tree Genetics & Genomes, 15(5), 1–13.

Ofori, A., & Padi, F. K. (2020). Reciprocal differences and combining ability for growth and yield components in cacao (Theobroma cacao L.): a case of recommended cacao varieties in Ghana. Euphytica, 216(12), 1–14.

Phillips-Mora, W., Castillo, J., Krauss, U., Rodríguez, E., & Wilkinson, M. J. (2005). Evaluation of cacao (Theobroma cacao) clones against seven Colombian isolates of Moniliophthora roreri from four pathogen genetic groups. Plant Pathology, 54(4), 483–490.

Pimenta, N.A.A., D. Laranjeira, J.L. Pires, and L.E.D.M. Newman (2018) Selection of cocoa progenies for resistance to witches’ broom. Tropi. Plant Pathol. 43: 381–388

Phillips-mora, W. (2015). Avances tecnológicos del Centro Agronómico Tropical de Investigación y Enseñanza (CATIE) en la cadena del cacao. 29–31.

Phillips-Mora W, Castillo J (1999) Artificial inoculations in cacao with the fungi Moniliophthora roreri (Cif. Par) Evans et al. and Phytophthora palmivora (Butl.) Butler. In: CATIE (ed) Actas. IV Semana CientíficaTurrialba. Logros de la investigacion para un nuevo milenio., Turrialba. Costa Rica. CATIE

Purwantara, A., McMahon, P., Susilo, A. W., Sukamto, S., Mulia, S., Saftar, A., … & Guest, D. (2015). Testing local cocoa selections in Sulawesi:(ii) resistance to stem canker and pod rot (black pod) caused by Phytophthora palmivora. Crop Protection, 77, 18–26.

Phillips-Mora, W., Cawich, J., Garnett, W., & Aime, M. C. (2006). The first report of frosty pod rot (moniliasis disease) caused by Moniliophthora roreri on cacao in Belize. Plant Pathology, 55(4), 584. 10.1111/j.1365-3059.2006.01378.x

Phillips-Mora, W., & Wilkinson, M. J. (2007). Frosty pod of cacao: A disease with a limited geographic range but unlimited potential for damage. Phytopathology, 97(12), 1644–1647. 10.1094/PHYTO-97-12-1644

Phillips-Mora, W., Arciniegas, A., Mata, A., & Motamayor, J. (2012). Catálogo de clones de cacao seleccionados por el CATIE para siembras comerciales. In Serie técnica - Manual técnica 105.

Phillips-Mora W, Mata-Quiros A, Arciniegas-Leal A (2017) Generation of cacao clones with durable resistance against moniliasis/frosty pod rot (Moniliophthora roreri). In: International symposium on cacao research, Lima, Peru

Phillips-Mora, W., & Cerda, R. (2009). Catálogo: enfermedades del cacao en Centroamérica. Centro Agronómico Tropical de Investigación y Enseñanza, CATIE. Turrialba, Costa, Rica.

Ploetz, R. C. (2007). Cacao diseases: Important threats to chocolate production worldwide. Phytopathology, 97(12), 1634–1639. 10.1094/PHYTO-97-12-1634

Rorer, J.B. (1918). Enfermedades y plagas del cacao en el Ecuador y m etodos mod-ernos apropiados al cultivo del cacao.Report, Agricultural Society of Ecuador,p. 80. Guayaquil: Imprenta do Diario Illustrado.

R Core Team, 2019. R: A Language and Environment for Statistical Computing.

Sánchez-Mora, F. D., Mariela Medina-Jara, S., Díaz-Coronel, G. T., Ramos-Remache, R. A., Vera-Chang, J. F., Vásquez-Morán, V. F., Troya-Mera, F. A., Garcés-Fiallos, F. R., & Onofre-Nodari, R. (2015). Potencial sanitario y productivo de 12 clones de cacao en ecuador. Revista Fitotecnia Mexicana, 38(3), 265–274. 10.35196/rfm.2015.3.265

Rodriguez-Medina, C., Arana, A. C., Sounigo, O., Argout, X., Alvarado, G. A., & Yockteng, R. (2019). Cacao breeding in Colombia, past, present, and future. Breeding Science, 69(3), 373–382.

Rodríguez-Polanco, E., Morales, J. G., Muñoz-Agudelo, M., Segura, J. D., & Carrero, M. L. (2020). Morphological, molecular, and pathogenic characterization of Phytophthora palmivora isolates causing black pod rot of cacao in Colombia. Spanish Journal of Agricultural Research, 18(2), e1003–e1003.

Sánchez, J., and González, L. (1989). Metodología para evaluar la susceptibilidad a moniliasis en cultivares de cacao (Theobroma cacao). https://repositorio.catie.ac.cr/xmlui/handle/11554/10738

Somarriba, E., & Hernández Rodríguez, M. (2013). La contribución del Proyecto Cacao Centroamérica al estímulo del sector cacaotero de Centroamérica. Agroforestería en las Américas; No. 49.

Somarriba, E., Cerda, R., Orozco, L., Cifuentes, M., Dávila, H., Espin, T., … & Deheuvels, O. (2013). Carbon stocks and cocoa yields in agroforestry systems of Central America. Agriculture, ecosystems & environment, 173, 46–57.

Sreenivas, M., Bhattacharjee, T., Sharangi, A. B., Maurya, P. K., Banerjee, S., Chatterjee, S., … & Chattopadhyay, A. (2020). Breeding chili pepper for simultaneous improvement in dry fruit yield, fruit quality, and leaf curl virus disease tolerance. International Journal of Vegetable Science, 26(5), 457–486.

Soria, J., & Enríquez, G. A. (1977). Genetic improvement for resistance to five cacao diseases.

Solís Bonilla, J.L., Zamarripa Colmenero, A., Pecina Quintero, V., Garrido Ramírez, E., & Hernández Gómez, E. (2015). Evaluación agronómica de híbridos de cacao (Theobroma cacao L.) para selección de alto rendimiento y resistencia en campo a moniliasis. Revista mexicana de ciencias agrícolas, 6(1), 71–82.

Toker, C., & y;anci, H. (2003). Selection of chickpea (Cicer arietinum L.) genotypes for resistance to Ascochyta blight [Ascochyta rabiei (Pass.) Labr.], yield and yield criteria. Turkish Journal of Agriculture and Forestry, 27(5), 277–283.

Thevenin, J. M., Rossi, V., Ducamp, M., Doare, F., Condina, V., & Lachenaud, P. (2012). Numerous clones are resistant to Phytophthora palmivora in the “Guiana” genetic group of Theobroma cacao L. PLoS ONE, 7(7). 10.1371/journal.pone.0040915

Torres de la Cruz, M., Ortiz García, C. F., Téliz Ortiz, D., Mora Aguilera, A., & Nava Díaz, C. (2011). Temporal progress and integrated management of frosty pod rot (Moniliophthora roreri) of cocoa in Tabasco, Mexico. Temporal Progress and Integrated Management of Frosty Pod Rot (Moniliophthora roreri) of Cocoa in Tabasco, Mexico, 31–36.

Vu VQ (2020). A biplot based on ggplot2. https://github.com/vqv/ggbiplot

Wei, T., Simko, V., Levy, M., Xie, Y., Jin, Y., & Zemla, J. (2017). Package ‘corrplot’. Statistician, 56(316), e24.

